# A Computational Model of the Effect of Short-Term Monocular Deprivation on Binocular Rivalry in the Context of Amblyopia

**DOI:** 10.1101/2021.07.31.453675

**Authors:** Norman Seeliger, Jochen Triesch

## Abstract

Treatments for amblyopia focus on vision therapy and patching of one eye. Predicting the success of these methods remains difficult, however. Recent research has used binocular rivalry to monitor visual cortical plasticity during occlusion therapy, leading to a successful prediction of the recovery rate of the amblyopic eye. The underlying mechanisms and their relation to neural homeostatic plasticity are not known. Here we propose a spiking neural network to explain the effect of short-term monocular deprivation on binocular rivalry. The model reproduces perceptual switches as observed experimentally. When one eye is occluded, inhibitory plasticity changes the balance between the eyes and leads to longer dominance periods for the eye that has been deprived. The model suggests that homeostatic inhibitory plasticity is a critical component of the observed effects and might play an important role in the recovery from amblyopia.

## 1 Introduction

Amblyopia (greek, meaning *dull* or *blunt sight*) is a developmental disorder of the visual system in which the brain and an eye are not working well together. For people suffering from amblyopia one eye — the then called amblyopic eye — shows little visual acuity as well as a decreased contrast and motion sensitivity [9]. Patients typically also suffer from poor stereo vision. Neither can be related to structural abnormalities nor can it be entirely recovered during adulthood. Amblyopia can be caused by, e.g., a muscle imbalance (strabismic amblyopia) as well as by a difference in sharpness of vision between the eyes (anisometropic amblyopia) during the first 3-5 years of life. Different causes lead to different characteristics of amblyopia, e.g., regarding visual acuity and contrast sensitivity [1]. Considering these causes and their consequences, the suppression of the amblyopic eye’s neural representation by the fellow eye seems to be the primary cause for a failing contribution of the amblyopic eye to vision. Indeed, it was found that for all major forms of amblyopia the degree of interocular, GABAergic suppression correlates with the depth of amblyopia [14, 21].

Under normal viewing conditions, the input to both eyes is nearly identical and just differs slightly due to different viewing angles. When both eyes are presented with non-matching input, however, most people report fluctuations between the perceptions of these inputs with the observer perceiving only one eye’s image at a time. This phenomenon is called binocular rivalry. The fluctuations in perception during binocular rivalry are stochastic and show a mean duration of about 2 seconds [4]. They are thought to arise due to a competition between neural populations representing the two different percepts, which is mediated by mutual inhibition. Recent computational models of binocular rivalry focused on addressing this mutual inhibition with alternations in perception being allowed due to adaptation [13][5] and/or noise in the network [20]. Since amblyopia and binocular rivalry both rely on competition between the eyes, they may be related at a mechanistic level.

A recovery from amblyopia can be achieved best during childhood, e.g., by the most common treatment incorporating eye patches during an occlusion therapy. Here, the strong eye is occluded for several hours per day over multiple months. Recently, Lunghi and others combined the patching therapy with a novel measure for neuronal plasticity which incorporates binocular dynamics [15–17]. They first showed that after a short-term monocular deprivation of 150 minutes, the occluded eye dominates perception under binocular rivalry in healthy adults [15]. The strength of this effect correlated with decreased GABA levels after patching [16], diminished over time, and went back to a normal state after 90 minutes. In a following study, they found that this effect is also present for the patched fellow eye of amblyopes undergoing standard occlusion therapy [17]. Moreover, the recovery rate from amblyopia could be predicted: the stronger the dynamics are altered after the patching, the more the amblyopic eye could recover after months of treatment. Zhou et. al. [27] added to this finding by successfully applying an inverse occlusion therapy in adult amblyopes for which, again, the binocular balance was the key aspect for recovery.

The physiological mechanisms responsible for the observed effects are still unknown, however. Therefore, the current work aims to provide a better understanding of potential mechanisms leading to the effects described above. We hypothesized that inhibitory plasticity may be the central mechanism giving rise to the observed effects. To test the plausibility of this hypothesis, we first created a spiking neural network model that produced alternations in dominance between two competing neuron groups, which were stimulated simultaneously. Second, one of the groups was deprived of its normal input for a certain amount of time. Inhibitory plasticity altered the rivalry dynamics during deprivation and lead to longer dominance durations for the previously deprived population. This effect was accompanied by a temporary reduction in GABA-levels, as observed experimentally. Based on these findings, we conclude that inhibitory plasticity is a plausible explanation for the observed experimental findings.

## 2 Methods

The current model aims to represent a part of layer IV of the primary visual cortex. It incorporates pyramidal neurons (excitatory, regular-spiking) and parvalbumin neurons (inhibitory, fast-spiking) in two interconnected layers. The excitatory neuron layer consists of 200 neurons and the inhibitory layer of 50 neurons to maintain ratios between these two groups as seen in experiments. The general structure of the model is displayed in Figure 1.

**Fig. 1.**
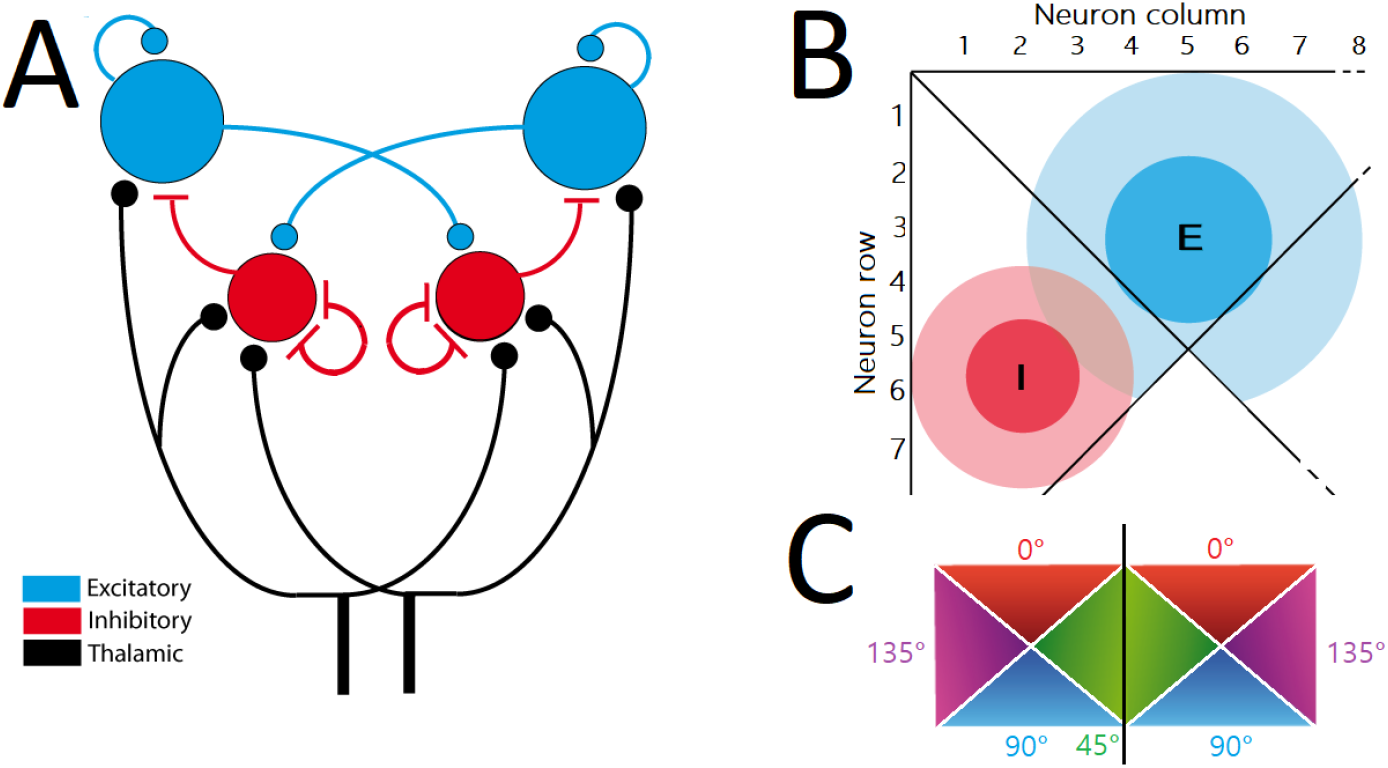
Model architecture and connectivity. **A)** Modeled layer IV with exc. (blue) and inh. populations (red) as well as thalamic drive (solid black). Excitatory synapses are displayed as bulbs, inhibitory synapses as lines. **B)** Connectivity within one ocular dominance column following a Gaussian probability function (exc: *σ* of 3, inh: *σ* = 2). Lines separate regions of exc. neurons that respond to the same orientation. The second column is placed adjacent (not shown) and shows the same connectivity. **C)** Pinwheel architecture of the model. Black line separates ocular dominance columns.

Each layer consists of two ocular dominance columns, which incorporate a pinwheel structure, once clockwise and once counter-clockwise, which indicate neuron populations with a varying orientation preference. For each dominance column, 100 exc. and 25 inh. neurons are divided into 4 groups which then share the same orientation preference (e.g. exc. neurons 1-25: left eye, preference of 0 degrees).

Neurons in the model are of the standard leaky-integrate-and-fire type. They include a slow hyperpolarizing current to model neuronal spike rate adaptation and white Gaussian membrane noise that is added to the membrane voltage. The general formulation is adapted from [6]. When the membrane potential of a neuron reaches a threshold (mean value *—*57.3 mV for exc. neurons, *−*58.0 mV for inhibitory neurons, both with a standard deviation of 0.1 mV), a spike is generated and the neuron’s membrane potential is set back to *−*70.0 mV for excitatory neurons and *−*60.0 mV for inhibitory neurons.

Synapses are modeled as conductance-based. Upon arrival of a presynaptic spike, the synapse’s conductance is increased by the synaptic weight *w* and then decreases exponentially with a time constant specific to the synapse type (*τ*_AMPA_ = 3 ms, *τ*_NMDA_ = 80 ms, 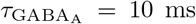). The different synapse types are used to model external stimulation by the thalamus (AMPA-mediated), recurrent excitation provided by adjacent excitatory neurons and not explicitly modeled layer V/VI neurons alike (AMPA- and NMDA-mediated) as well as inhibition from inhibitory neurons (GABA_A_-mediated).

The spike rate adaptation mechanism works similarly. Upon each postsynaptic spike, the conductance is increased by a parameter *w*_SRA_ and then decays exponentially with the time constant *τ*_SRA_ = 996 ms.

Considering plasticity in the model, STDP was implemented only at inhibitory to excitatory connections for which a symmetric STDP window has been assumed with a pairing of pre- and postsynaptic spikes leading to potentiation independent of the relative order of the events:

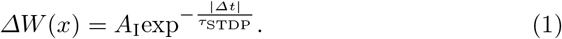

Here, *Δt* denotes the time between pre- and postsynaptic spikes, *τ*_STDP_ is the time constant for the STDP and *A*_I_ is a constant for the maximum possible weight change. An online implementation of a nearest-neighbor STDP rule is used with every spike leading to a trace, which is evaluated once a spike of the corresponding pre- or postsynaptic neuron occurs. For homeostatic purposes, an *α*-parameter is also introduced which reduces the synaptic weight by a small amount *α* for every presynaptic spike of an inhibitory neuron and which is independent of spike times of the corresponding postsynaptic excitatory neuron [25].

The global connectivity of the network promotes inhibition between groups in different ocular dominance columns which are driven by different sensory stimuli. This is in line with experimental findings showing that inhibition is prominent from one ocular dominance column towards the other [10]. This inhibition is mediated by long-range excitatory connections from excitatory neurons of one eye’s dominance column towards inhibitory neurons of the other eye’s dominance column. For the local connectivity within one ocular dominance column, every neuron pair has a chance to become connected that drops of with increasing distance following a Gaussian probability function (exc: *σ* of 3, inh: *σ* = 2; see Figure 1B). Thus, the longest axons stem from excitatory rather than inhibitory neurons [19]. Connectivity parameters were adapted from Ahmed et. al [2] for the relative contributions of thalamic and cortical projection ratios and from Potjans et. al [23] for cortico-cortical connection ratios. Axonal delays are heterogeneous (exc. to exc. 1.5 ms, inh. to exc. 0.5 ms, exc. to inh. 1.0 ms and inh. to inh. 1.0 ms).

Every neuron is stimulated via 10 spike trains (40 Hz, interocular correlation of 0.08 and intraocular correlation of 0.25). The same input is also provided to the inhibitory populations which target the excitatory populations of both eyes (see Fig. 1). During the simulated patching, the input towards one eye is set to 10 Hz and is uncorrelated while the input to the other eye is left unchanged.

For the computation of dominance durations, mean firing rates are calculated for the two groups belonging to the eyes using a rectangular sliding window with a width of 300 ms. The population which is more active than the other population is labeled as *dominant* at this moment. Switches in dominance are indicated when the suppressed population becomes at least twice as active as the formerly active population. Periods with mixed perceptions and transitions between dominance durations of competing groups are not treated differently for the current analysis. For the comparison of distributions of dominance durations, a Kolmogorov-Smirnow test is used for the dominance durations of one eye before and after the patching.

The model is implemented using Brain2 [24]. The program code is available at GitHub under https://bit.ly/38jZQgX and licensed under GNU General Public License v3.0.

## 3 Results

### 3.1 General Network Behavior

An example of the network’s activity is given in Figure 2. Here, the excitatory populations representing the left (bottom row) and right (top row) eye compete for dominance when neurons that prefer orthogonal orientations are stimulated simultaneously. These populations show varying dominance durations with a mean slightly above 2 seconds as seen in experiments [4] (Figure 3). The inhibitory populations show the inverse activity (data not shown).

**Fig. 2.**
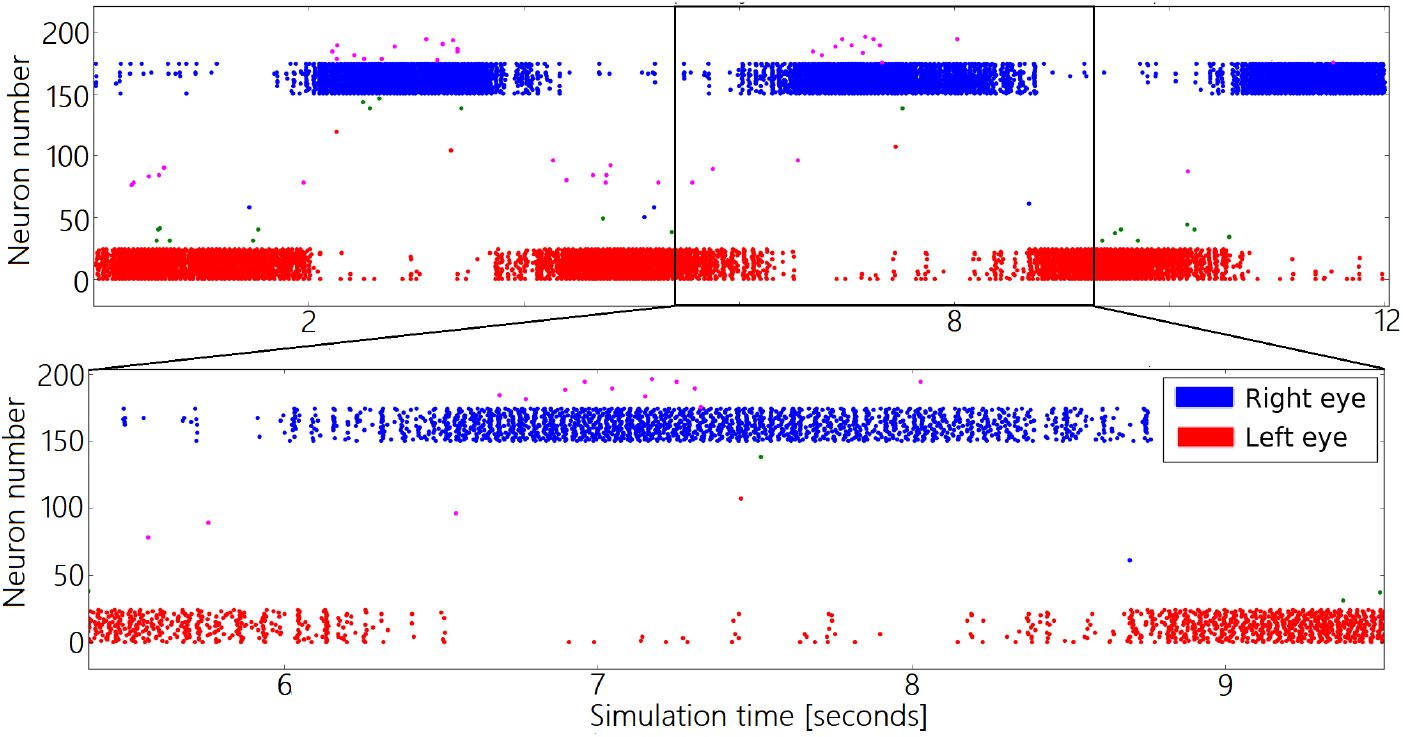
Rasterplots of network activity. Example of the network activity under basal rivaling conditions. The stimulated populations are left eye, 0 deg. (red) and right eye, 90 deg. (blue). Other colors correspond to other eye, orientation combinations.

**Fig. 3.**
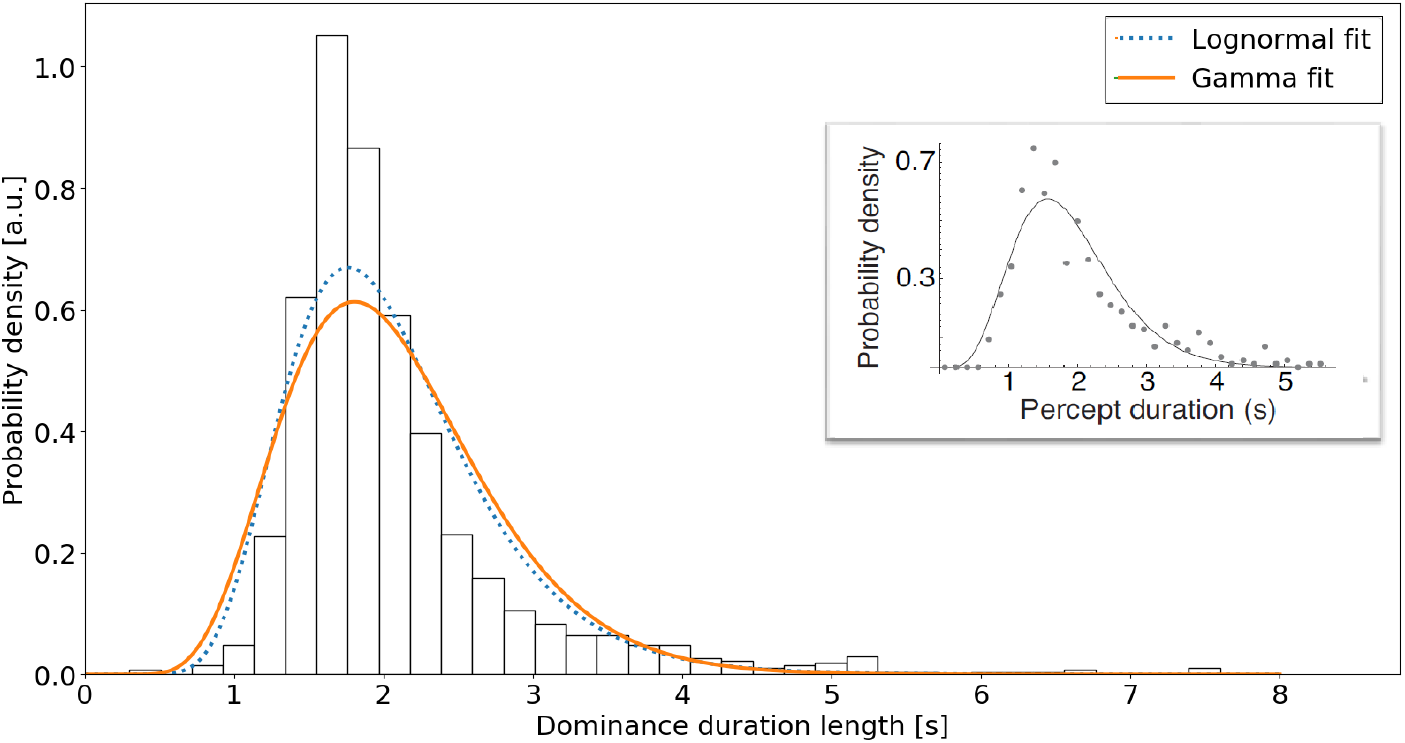
Dominance duration distribution. Distribution of dominance durations for the model together with a gamma fit and a lognormal fit. Inset: Perception duration lengths found in experiments with a fitted gamma distribution (see [4]).

### 3.2 Effect of Patching

We modeled the patching procedure used in experiments. Participants typically watch a movie containing a wide range visual stimuli or are free to perform their normal everyday activities while the patch is applied. The stimulus before and after the patching depends on the particular experiment (e.g. unrestricted input or specific tasks). For the purpose of our model, we provide the rivaling input for the entire time. Dominance durations are calculated for the first and last third, inhibitory weights are recorded for the entire time. To save simulation time, we accelerated the effect of plasticity to reduce the time required to simulate the occlusion to 100 seconds. Figure 4 shows a representative example of the effect of the simulated patching on the network dynamics. In part (A), the mean of the inhibitory weights targeting the corresponding excitatory populations is shown with the green/red vertical lines indicating the start/end of the occlusion. Before the patch is applied, the inhibitory weights towards both populations slightly diverge since one population is slightly stronger than the other due to random factors in the initialization of the network. As soon as the patching starts, however, the mean inhibitory weight towards the occluded eye drops substantially while the mean inhibitory weight towards the open eye shows an increase. When the patching stops, the reverse is visible: the strengths of inhibitory weights towards the formerly occluded eye rise again to the level of the unoccluded eye. The mean inhibitory weight towards the unoccluded eye starts to decrease. Both means seem to approach each other in a time frame comparable to the patching duration.

**Fig. 4.**
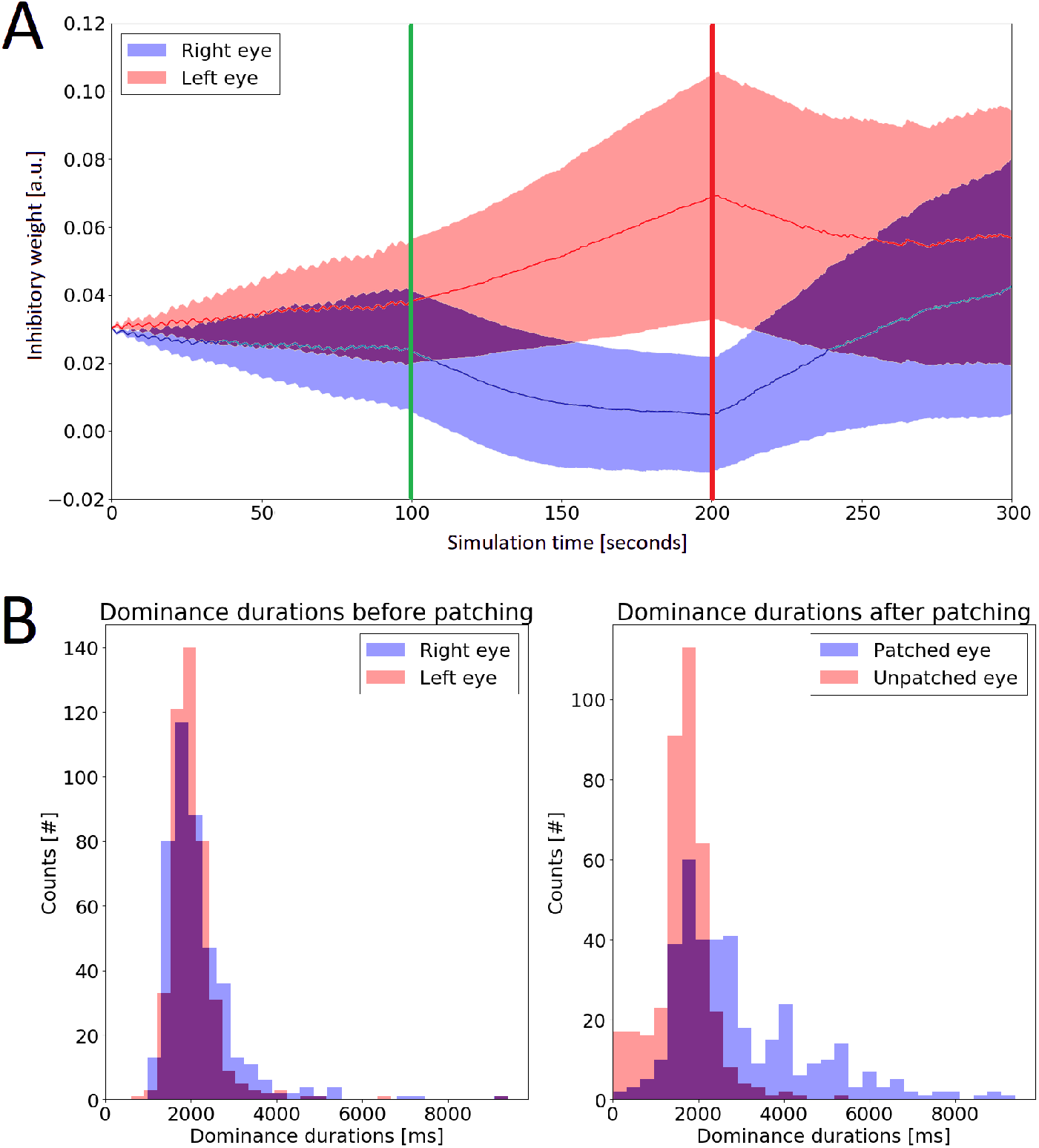
Effect of patching. **A)** Evolution of the inhibitory weight (mean and standard deviation) targeting excitatory neurons of the open (top) and occluded eye (bottom). Vertical green and red lines indicate, respectively, the start and end of the occlusion. One exemplary weight evolution was chosen. **B)** Dominance durations before (left) and after (right) the patching (combined data from 18 simulations).

The impact of these altered weights on rivaling dynamics can be seen in Figure 4B (data from 18 different simulations). Before the patching, every eye shows average dominance durations of roughly the same length (left: 2069 ms; right: 2164 ms). Afterwards, this behavior is dramatically altered. The formerly occluded population can be active for very long time periods and now possesses a significantly prolonged mean dominance duration of 3212 ms (KS-Test: *p <* 0.001). The eye which stayed open during the patching has its mean dominance duration significantly reduced to 1667 ms (KS-Test: *p <* 0.001). The change in dominance duration length for the occluded eye is stronger than the change for the eye which remained open. Thus, the model reproduces the changes in dominance durations after patching observed experimentally.

## 4 Discussion

We aimed to create a model that captures and explains the experimental effects observed by Lunghi and others [15–18]. The main focus lied in creating a spiking neural network model with inhibitory plasticity to explain how the dynamics of binocular rivalry are altered in response to monocular deprivation. Our network captures these effects and demonstrates how the occlusion of one eye can lead to a temporary relief of the corresponding part of the primary visual cortex from inhibition. This then allows for a re-balancing of the total network undisturbed from a potential suppression by the other eye.

In the model and with the occlusion of one eye, the input towards inhibitory neurons targeting the corresponding excitatory population is only partially decreased due to the drive coming from the eye which remains open. This presynaptic inhibitory activity, however, has little to no postsynaptic excitatory activivity with which the spikes could be correlated. Thus, the inhibitory weight of these synapses decreases over time during the occlusion and only starts to regain strength after the patch is lifted. At this third stage, the chance of correlated activity is elevated which leads to a strong potentiation of the targeting inhibitory weight. A similar, but inverted, effect can be seen for the eye which remains open: during the occlusion, the inhibitory neurons for the open eye gain nearly all of their input from the same neurons which drive the excitatory population. Thus, the activity is highly correlated and leads to a potentiation of the average inhibitory weight. Interestingly, we also found a similar impact of the patching in a variant of this network without feed-forward inhibition (data not shown). In this variant, the inhibition towards the open eye remained stable but the occluded eye experiences a decreased inhibition. Both versions of the model lead to the occluded population dominating after the patching under rivaling conditions. This effect is also robust with respect to the condition of the network at the moment of patching — since the strength of the eyes differs in healthy people in general and in amblyopic people in particular, completely balanced conditions would be a rather unrealistic setting. The reaction of the network to the occlusion directly results from the used parameters (e.g. for the STDP) and thus, the ability of the network to show plastic changes due to this treatment could be linked to the overall plasticity of the brain. This could explain why Lunghi et al. could predict the recovery rate from amblyopia based on the impact of short-term occlusion on binocular rivalry.

The network also agrees with details of the results by Lunghi et. al [15]. The increase in dominance duration length of the formerly occluded eye exceeds the decrease for the eye which remained open. Also, the time constants seem to fit: in the experiment, the patching had shown an impact for a duration which is slightly shorter than the time the patching was performed. A similar time frame can be seen in the model.

Also, our model is robust to various design choices. In the network, excitatory neurons only connect to excitatory neurons with a rather similar orientation preference (up to 45 degrees) while avoiding excitatory neurons with opposite preferences. However, an architecture where excitatory neurons only connect to neurons showing the same preference would also be plausible. This approach leads to similar results in the current model. The same is true for a variant of the model where projections crossing the ocular dominance column border are not provided by excitatory neurons towards inhibitory populations, but by inhibitory populations targeting the other eye. This architecture also yielded comparable results.

But what might be the implications of reduced inhibition in the brain? Parvalbumin-positive (PV+) inhibitory neurons play an important role in guiding cortical plasticity, with the maturation of these neurons marking the onset of critical periods, e.g., in the visual cortex. Reopened periods for ocular dominance plasticity later in life are, however, achieved through reduced inhibition. This is shown for example by Kuhlmann et al. [12], who re-enabled juvenile-like plasticity in the visual cortex by artificially inhibiting the activity of PV+-neurons. This then allowed excitatory neurons to become plastic again. Barnesd et al. [3] added the finding that the recovery of neurons responding preferentially to a patched eye depends on the amount of correlated activity, which matches the findings of our model. With regards to amblyopia, there also is recent computational evidence highlighting the importance of cortical plasticity for a potential recovery [8]. There are different possible mechanisms of how PV+ neurons can guide cortical plasticity, one of which is the strong effect of perisomatic inhibition onto backpropagating action potentials and the temporal window in which arriving inputs can sum up and provoke a response of the target neuron. A release of that inhibition together with increased excitability can help these otherwise suppressed neurons to compete, which is important to consider for strongly suppressed populations that represent an amblyopic eye. Another interesting aspect is a possible effect of PV+ neurons on the tPA enzyme (tissue plasminogen activator), which is more active following monocular deprivation and supports pruning mechanisms, which are important for ocular dominance plasticity [22].

Most of these aspects, however, take place in higher layers such as layer II/III. Nevertheless, a key role of parvalbumin-positive neurons and their plasticity was made clear by the studies mentioned above. PV+ neurons also receive potent thalamic input [11], show a specific degree of orientation tuning [26], and change their levels of activity under locomotion [7]. The latter point is consistent with [18] showing increased effects of the patching paradigm when combined with physical exercise. Therefore, inhibitory neurons in general and parvalbumin neurons, in particular, could be a key player in plastic changes also in layer IV during ocular dominance alterations and an important mediator of a possible recovery from amblyopia.

## Acknowledgments

This work was supported by an ERA-NET NEURON grant (JTC2015), the German Ministry for Education and Research (BMBF, grant number 01EW1603A). JT acknowledges support from the Johanna Quandt foundation. The authors thank Dr. Florence Kleberg for her input and contributing ideas as well as Samuel Eckmann and Lukas Klimmasch for support and fruitful discussions.

